# Impact of a rapid decline in malaria transmission on antimalarial IgG subclasses and avidity

**DOI:** 10.1101/2020.06.26.173005

**Authors:** Isaac Ssewanyana, John Rek, Isabel Rodriguez, Lindsey Wu, Emmanuel Arinaitwe, Joaniter I Nankabirwa, James G Beeson, Harriet Mayanja-Kizza, Philip J Rosenthal, Grant Dorsey, Moses Kamya, Chris Drakeley, Bryan Greenhouse, Kevin K.A.Tetteh

## Abstract

Understanding how immunity to malaria is affected by declining transmission is important to aid vaccine design and understand disease resurgence. Both IgG subclasses and avidity of antigen-specific responses are important components of an effective immune response.

Using a multiplex bead array assay, we measured the total IgG, IgG subclasses, and avidity profiles of responses to 18 *P. falciparum* blood stage antigens in samples from 160 Ugandans collected at 2 time points during high malaria transmission and 2 time points following a dramatic reduction in transmission.

Results demonstrated that, for the antigens tested, (i) the rate of decay of total IgG following infection declined with age and was driven consistently by the decrease in IgG3 and occasionally the decrease in IgG1; (ii) the proportion of IgG3 relative to IgG1 in the absence of infection increased with age; (iii) the increase in avidity index (the strength of association between the antibody and antigen) following infection was largely due to a rapid loss of non-avid compared to avid total IgG; and (iv) both avid and non-avid total IgG in the absence of infection increased with age.

Further studies are required to understand the functional differences between IgG1 and IgG3 in order to determine their contribution to the longevity of protective immunity to malaria. Measuring changes in antibody avidity may be a better approach of detecting affinity maturation compared to avidity index due to the differential expansion and contraction of high and low avidity total IgG.

## Introduction

IgG is an important component of immunity to malaria, with function defined by the specific recognition of antigens, and tropism of the constant region (Fc) for variant Fc receptors (1). Avidity, the sum of binding affinities between antibodies and antigenic epitopes, may reflect the functional quality of the antibody variable region (2,3). IgG subclasses IgG1 - IgG4 have differences in the Fc region that affect their affinity to variants of the Fcγ receptors, influencing their effector function, longevity, and ability to cross the placental barrier (4,5). IgG subclass switching and affinity maturation(6,7), are important components that dictate the quality of immunity to malaria (8–10).

Antibody levels and breadth of response to specific *P. falciparum* antigen targets generally diminish in the absence of re-infection, which is thought to contribute to loss of immunity (11–16). However, there are differences in the rate of decay of antibodies to different antigens (17,18). Antibody responses to malaria are predominantly cytophilic (IgG1 and IgG3) and have been shown to mediate effector mechanisms that inhibit of parasite growth (19,20), promote opsonic phagocytosis (21)) and complement fixation (22,23). Epidemiological studies have demonstrated associations between IgG1 and IgG3 targeting various *P. falciparum* antigens and different manifestations of immunity, including reductions in the risk of infection, parasite density, clinical disease, and disease severity (15,20,24,25). In general, these associations were stronger for IgG3 compared to IgG1 (26–28). Furthermore, in-vitro functional assays have implicated interference by IgG2 and IgG4 in the opsonizing function of the cytophilic IgG1 and IgG3 antibodies in competition assays (29,30).

Previous studies have shown differences in class switch bias profiles driven by different *P. falciparum* antigens (31–34). Other factors such as age and cumulative exposure are also thought to influence subclass switching (35). Therefore, the relative composition of the subclasses may influence the functional relevance of antibodies in antimalarial immunity. Antibody avidity is a correlate for immune memory and protection in some infections (36–42). In malaria, studies have described an increase in affinity following resolution of clinical malaria (43), higher avidity in those with reduced risk of complicated malaria (44,45), higher avidity in clinically immune compared to non-immune populations (43), and an association between higher avidity and reduced risk of placental malaria (46). Surprisingly, a prior study by our group demonstrated that avidity to the *P. falciparum* antigens AMA-1 and MSP1-19 was inversely related to transmission intensity at 3 sites in Uganda (47). This seemingly counterintuitive result prompted us to more measure avidity to broader array of antigens and to explicitly evaluate changes over time in individuals living in a setting of changing transmission intensity.

Few studies have combined the evaluation of IgG subclasses and avidity, and responses to only a limited number of *P. falciparum* antigens have been studied. We also do not fully understand the natural history of antibody waning in the absence of infection that may be important for naturally acquired immunity and its longevity. Understanding how naturally acquired immunity is maintained is important to identify populations at risk in settings with declining malaria transmission and to better inform malaria vaccine design.

In this study, we measured antibody levels to total IgG, IgG subclasses and avidity index to 18 *P. falciparum* blood stage antigens in the same individuals before and after decreases in malaria transmission (>90%) due to indoor residual spraying of insecticide (IRS). We compared the net changes in antibody responses during this period to gain a broader insight into how IgG subclasses and avidity index are acquired with age, and how they influence a reduction in IgG levels in the absence of infection.

## Methods

### Study population

Study participants were part of a cohort in Nagongera, eastern Uganda described in detail elsewhere (48). In 2011 Nagongera had one of the highest malaria burdens in the region with an entomological inoculation rate (EIR) of 215 infectious bites per person per year (49). Malaria control interventions included the use of insecticide treated nets (ITN), malaria case management with artemisinin-based therapies and intermittent presumptive treatment during pregnancy with sulfadoxine-pyrimethamine as per the Ugandan National Malaria Control Program policy. Between December 2014 and February 2015, IRS with the carbamate bendiocarb was introduced for the first time, followed by additional rounds ~ every 6 months thereafter. This intervention led to a dramatic decrease in the burden of malaria (Figure1).

**Figure 1.**
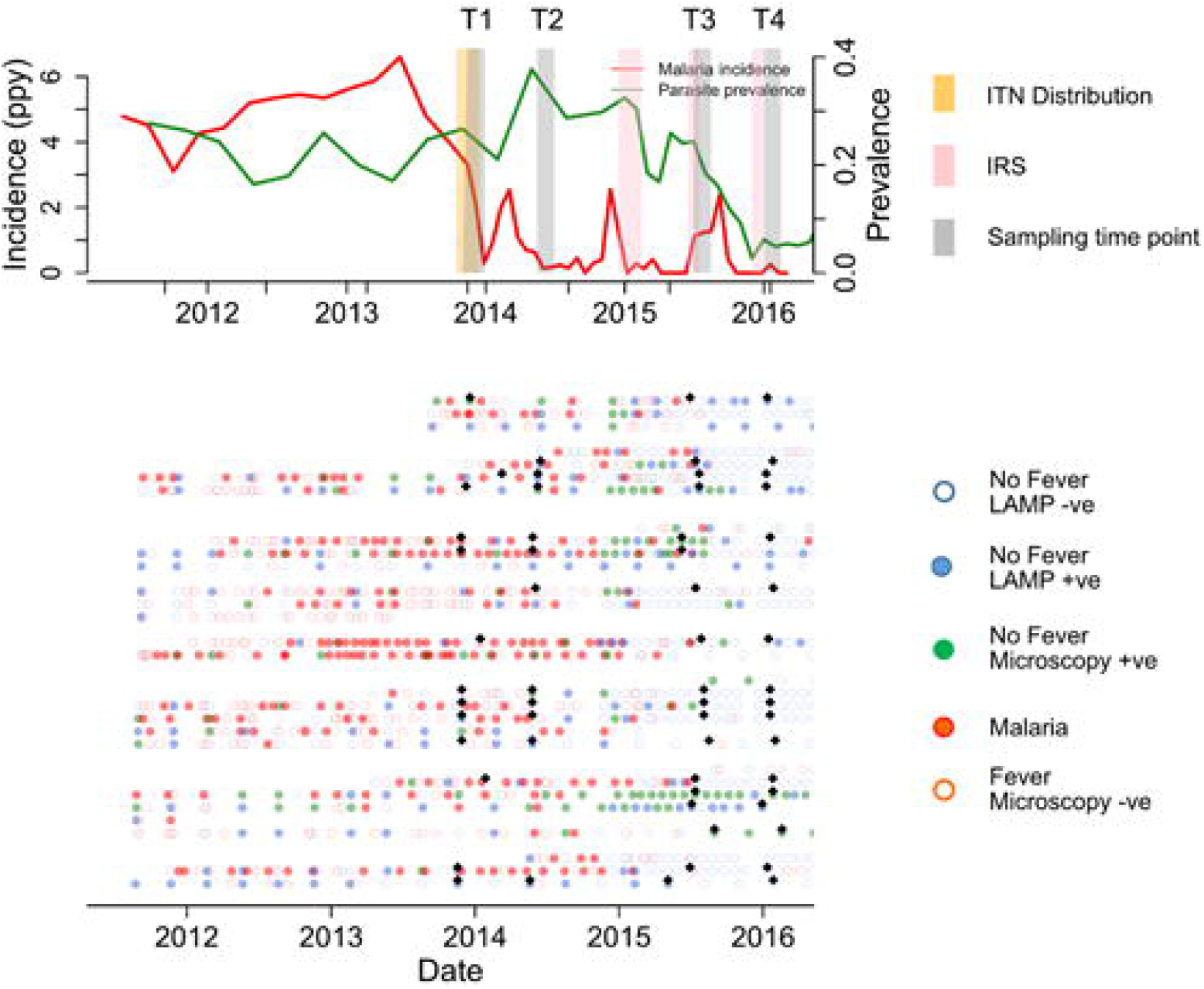
Summary of cohort at population and individual level. Upper panel shows malaria incidence, parasite rate, ITN distribution (yellow bar), and IRS spraying schedules (pink bars) in the full cohort. Lower panels are individual malaria status and 160 study sampling time points (black diamonds), at T1, T2, T3 and T4 (6, 12 months before first IRS and 6, 12 months after first IRS respectively). Participants were actively followed initially at 3 months interval and in Oct 2016 increased to 1 month. Patent by microscopy and sub-patent by LAMP Infections are represented by green filled and blue filled circles respectively and malaria fevers (red filled). Red filled, and red open represent malaria and non-malaria fever respectively. Green open, blue open circles represent microscopy and LAMP negative respectively.

Plasma samples were selected from study participants (n=160) and classified into 3 age groups; 1 - 4 (n=40), 5 - 11 (n=92) and >18 years (n=28). Time points were selected at 12 and 6 months pre-IRS (T1 and T2, respectively), and at 6 and 12 months post-IRS (T3 and T4, respectively). Study participants were provided with ITN at enrollment and monitored for parasite prevalence and malaria incidence (48). Malaria cases were identified via passive surveillance. *P. falciparum* infection was identified by active surveillance at 1 - 3 monthly intervals using both microscopy and LAMP (50).

### Ethical approval

Ethical approval was obtained from the Makerere University School of Medicine Research and Ethics Committee (REC REF 2011-203), the Uganda National Council for Science and Technology (HS 1074), the LSHTM ethics committee (reference # 6012), and the University of California, San Francisco Committee on Human Research (reference 027911).

Written informed consent was obtained from a parents or guardian of each child enrolled in the study and also from participating adults.

### *P. falciparum* antigens

A total of 18 recombinant *P. falciparum* blood stage antigens were assessed in addition to tetanus toxoid (TT) (National Institute of Biological Service and Control) as a non-malaria control. The *P. falciparum* antigens broadly fell into three categories; (i) infected red blood cell, (ii) merozoite apical organelle, and (iii) merozoite surface antigen associated proteins. The recombinant antigens were expressed in *Escherichia coli* either as glutathione S-transferase (GST) fusion proteins or as histidine tagged constructs (supplementary table1), with the exception of AMA1(51) which was expressed in *Pichia pastoris* and Rh5.1 which was expressed in HEK293 mammalian cells (52).

### Multiplex bead array assay to measure total IgG and IgG1 - 4

Total IgG responses to 18 *P. falciparum* blood stage antigens and TT were assayed in plasma at 1/1000 dilution using a multiplex bead array assay. Antigens were coupled to magnetic MagPlex microsphere beads (Luminex Corp, Austin, Texas) and assayed as previously described (53). Briefly, 50μl of the pooled bead suspension was added to each well, washed and incubated with 50μl of 1/1000 test plasma or control sample for 1.5hr at room temperature. The plates were washed and goat anti-human IgG rPE labelled antibody (Jackson Immuno Research Laboratories) was added and incubated for 1.5hr on a shaking platform. The plates were washed, 1×PBS added and read on a MagPix machine (Luminex Corp, Austin, Texas). The results were expressed as median fluorescent intensity (MFI). A standard curve using a hyper immune plasma PRISM pool (PP) 4 was included on each plate to normalize for plate to plate variations. The blank well MFI was deducted from each well to determine the net MFI.

The assay to measure total IgG was modified to measure IgG subclass responses at 1/100 including a two-step biotinylated anti-human IgG subclass (mouse anti-human IgG1: HP6069, IgG2: HP6002, IgG3: HP6050 IgG4: HP6023, Thermo fisher Scientific, UK) with a streptavidin-Phycoerythrin tertiary (Thermo fisher Scientific, UK), similar to what was previously described (54,55). The mouse anti-human secondary antibodies were used at 1/400, 1/400, 1/1,000, and 1/400 for IgG1 - 4, respectively. The results were expressed as MFI was after subtracting the blank well.

### Multiplex bead array assay to measure IgG avidity index

To measure the avidity index, the total IgG assay was modified to include a Guanidine Hydrochloride (GuHCl) step to dissociate antibodies with low avidity. Briefly, 50μl GuHCl was added to one of the duplicate wells for each sample, after the plasma incubation step. PBS was added to the second well. After washing R-Phycoerythrin-conjugated AffiniPure F (ab’) 2 Goat anti-human IgG was added and processed as per the standard MagPix assay described above. The signal detected in the well treated with GuHCl represented high avidity antibodies. Avidity index was calculated as a percentage of the high avidity antibodies of the total antibody as shown below

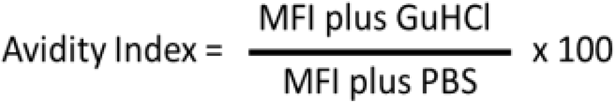

### Statistical analysis

The net changes in total IgG, IgG1 - 4, avidity index (AI) and the avid IgG levels (the antibody remaining bound after GuHCl treatment) between the peak and lowest malaria transmission were derived from a paired difference between log_10_ transformed MFI and AI at T2 and T4 respectively.

Associations between antibody levels (log_10_ transformed MFI) for total IgG, IgG1 - 4, avidity index and avid IgG with 90 days since last infection were assessed using the generalized estimation equation (GEE) to allow for repeated measures per individual including all 4 time points. Infections were defined by microscopy or loop mediated isothermal amplification (LAMP) for the smear negative slides.

Associations between total IgG, IgG1 - 4, avidity index and avidity index with age was assessed using a GEE model, adjusted for days since infection. Age groups 1 - 4 years was as the reference and compared to age groups 5 -11 and 18 years.

Median (and interquartile range) differences for total IgG, IgG1 - 4, AI and avid IgG levels between age categories where compared in a one-way ANOVA, not assuming normal distribution (using nonparametric, Kruskal - Wallis test) with pairwise comparisons between all age groups.

Analysis were performed and figures generated using Stata version 14 (StataCorp LLC, USA), GraphPad Prism version 7 (Graphpad Software Inc, USA) and R-studio (RStudio, Inc, USA).

## Results

### Decline in IgG levels against *P. falciparum* blood stage antigens with decreasing transmission

IgG responses were detected to all 18 *P. falciparum* antigens evaluated, with median responses higher for IgG3 compared to IgG1 to all antigens except HSP40 ag1 and MSP1-19, for which responses were similar, and to AMA-1 and Rh2_2030 which were higher for IgG1 compared to IgG3 (Supplementary Figure 1). To determine if the reduction in transmission in Nagongera was associated with antibody responses, we assessed the difference in antibody levels between T2 (6 months pre-IRS; peak transmission) and T4 (12 months post IRS; lowest transmission) (Figure 2 and Table 1). IgG levels (Log_10_MFI) were reduced for most of the *P. falciparum* antigens between the times of peak malaria transmission (T2) and lowest transmission (T4) (Figure 2A). A similar reduction was observed for almost all IgG3 responses, but this was not seen for IgG1, 2 or 4 responses. The reductions in the total IgG and IgG3 responses were similar for all the antigens tested. The largest reduction in total IgG was observed for Etramp5 ag1 and the smallest for AMA-1. The largest reduction in IgG3 was also observed for Etramp5 ag1 and the smallest for Rh5.1. IgG responses to tetanus toxoid, a non-malaria antigen control, did not change markedly between the two time points, consistent with the notion that the observed decreases were malaria specific. Taken together, these results showed that median total IgG levels declined dramatically between T2 and T4, mostly driven by reductions in IgG3 levels.

**Figure 2.**
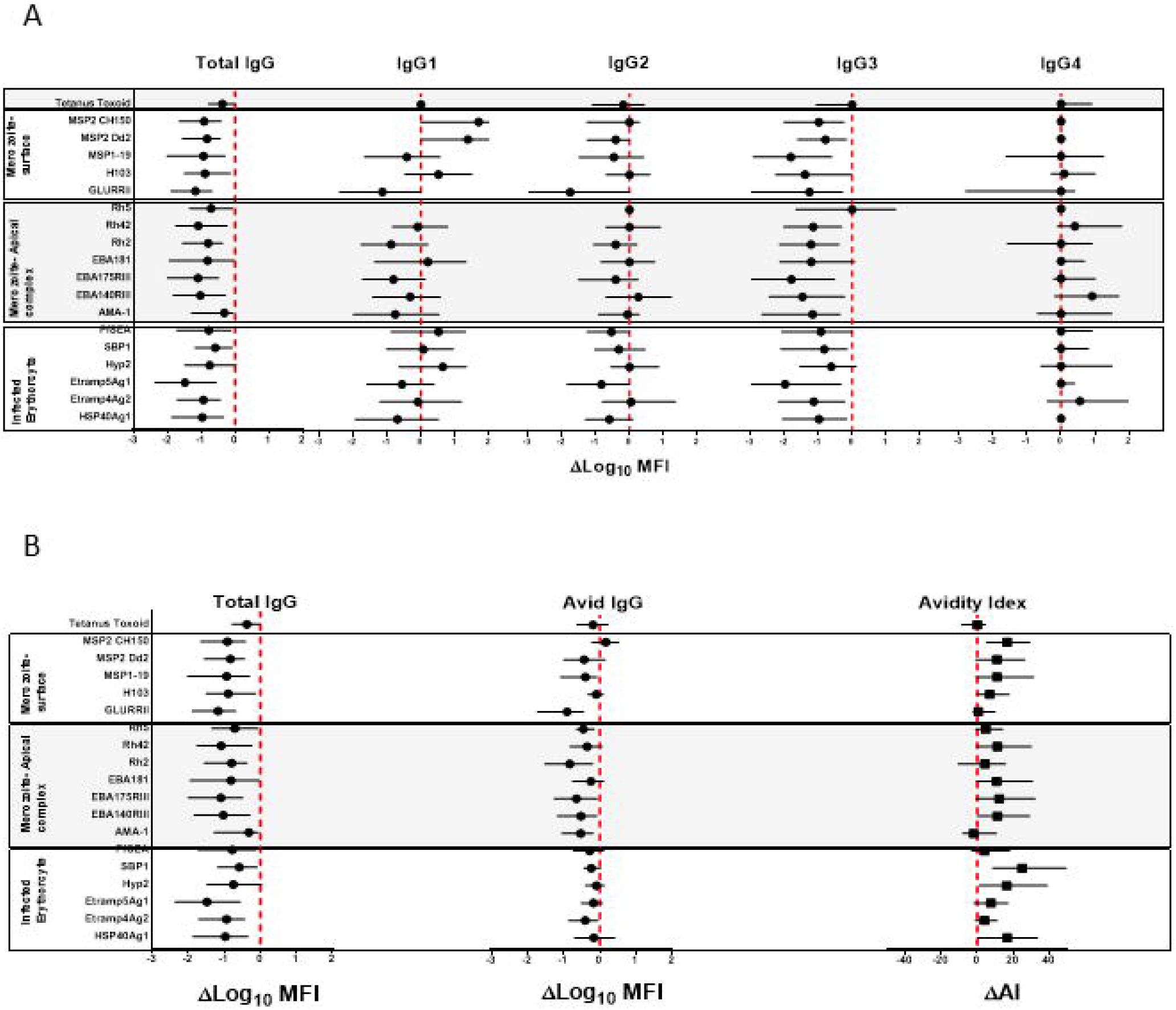
Net change in the median (IQR) of log_10_ MFI of total IgG, IgG1 – 4, avid IgG and avidity index between peak malaria transmission at T2 and near zero transmission at T4. (A) Total IgG and IgG1, IgG2, IgG3 and IgG4 (B) Total IgG and avid IgG and Avidity index. The dotted red line denotes zero; no net change in responses.

**Table 1.** Summary of population and malaria metrics. Time points were relative to the first round of IRS. T1=12, T2=6 months before IRS. T3=6, T2=12 months after IRS. Malaria was defined by fever and positive blood smear microscopy. Parasitemia status was defined by microscopy or LAMP positive at monthly interval

| Population and Malaria matrix | Age<br>(years) | Time point |  |  |  |
| --- | --- | --- | --- | --- | --- |
|  |  | T1 | T2 | T3 | T4 |
| Population | 1- 4 | 40 | 40 | 40 | 38 |
|  | 5 - 11 | 92 | 92 | 91 | 92 |
|  | >18 | 28 | 28 | 28 | 28 |
| Proportion with at least 1 clinical malaria episode in the last 90 days (%) | 1- 4 | 61 | 73 | 39 | 0 |
|  | 5 - 11 | 44 | 55 | 34 | 4 |
|  | >18 | 11 | 10 | 0 | 0 |
| Proportion with at least 1 parasitemia episode in the last 90 days (%) | 1- 4 | 76 | 80 | 58 | 14 |
|  | 5 - 11 | 95 | 89 | 85 | 45 |
|  | >18 | 58 | 70 | 45 | 14 |
| Mean days since parasitemia | 1- 4 | 51 | 40 | 91 | 240 |
|  | 5 - 11 | 9 | 23 | 50 | 146 |
|  | >18 | 77 | 84 | 186 | 322 |
| Proportion of months free of parasite in the last 6 months | 1- 4 | 0.42 | 0.35 | 0.62 | 0.83 |
|  | 5 - 11 | 0.27 | 0.25 | 0.41 | 0.62 |
|  | >18 | 0.48 | 0.53 | 0.71 | 0.85 |
| Proportion of months free of parasite in the last 12 months | 1- 4 | 0.45 | 0.40 | 0.51 | 0.75 |
|  | 5 - 11 | 0.29 | 0.29 | 0.37 | 0.51 |
|  | >18 | 0.49 | 0.51 | 0.64 | 0.78 |

### Increase in antibody avidity index to the *P. falciparum* blood stage antigens with decreasing transmission

To determine if the reduction in malaria transmission was associated with antibody avidity, we measured differences in the avidity index and the avid total IgG (i.e. antibodies remaining after the GuHCl antibody dissociation step), between T2 and T4. The median avidity index increased for most of the *P. falciparum* antigens between T2 and T4, with the exception of GLURP RII, AMA-1, Rh5.1 and Etramp5 ag1 (Figure 2B). The largest net increase in avidity index was observed for antigens SBP1 and HSP40 ag1. The high avidity antibodies showed a minor decrease in the antibody responses for most antigens (Figure 2B). For GLURP RII, Rh2_2030, EBA181 RIII-V, EBA175 RIII-V, EBA140 RIII-V and AMA-1, the magnitude of the decrease in the high avidity antibodies was less than that of total IgG. There were no consistent patterns across the different categories of the *P. falciparum* antigens. Avidity index and avid pool to tetanus toxoid did not change between T2 and T4, as expected, again indicating that the observations were specific for *P. falciparum*. Since the difference between total IgG and the avid pool was an indirect measure of low avidity antibodies, these results indicated that increased avidity index was driven by a preferential loss of the pool of antibodies with lower avidity following marked reduction in transmission.

### The decline in total IgG responses in the absence of *P. falciparum* infection was driven by preferential decay of IgG3

We hypothesized that population level changes in antibody responses associated with a decrease in malaria transmission were due to a waning of antibodies in the absence of blood stage infection. To test this hypothesis and measure the rate of change, we took advantage of frequent active and continuous passive surveillance for *P. falciparum* infection to relate antibodies at all 4 time points (T1 - T4) to the time since an individual’s last infection. Waning of antibodies was estimated using log-linear regression, i.e. assuming an exponential rate of decay and accounting for repeated measures using generalized estimating equations (GEE). Total IgG responses decreased following infection for all 18 *P. falciparum* antigens (Figure 3A). A higher magnitude of decrease following infection was observed in IgG3 for all antigens except Rh5.1 IgG1 responses were more heterogeneous, showing decreased, unchanged, or increased levels following infection (Figure 3A). IgG2 responses showed little or no change following infection. IgG4 responses reduced minimally for all antigens except GLURP RII and Etramp5 ag1. There was no change in tetanus toxoid responses following *P. falciparum* infection, confirming that these results were malaria specific. Taken together, the results demonstrated that waning total IgG responses following infection were consistently driven by decreases in IgG3 and occasionally by decreases in IgG1 responses.

**Figure 3.**
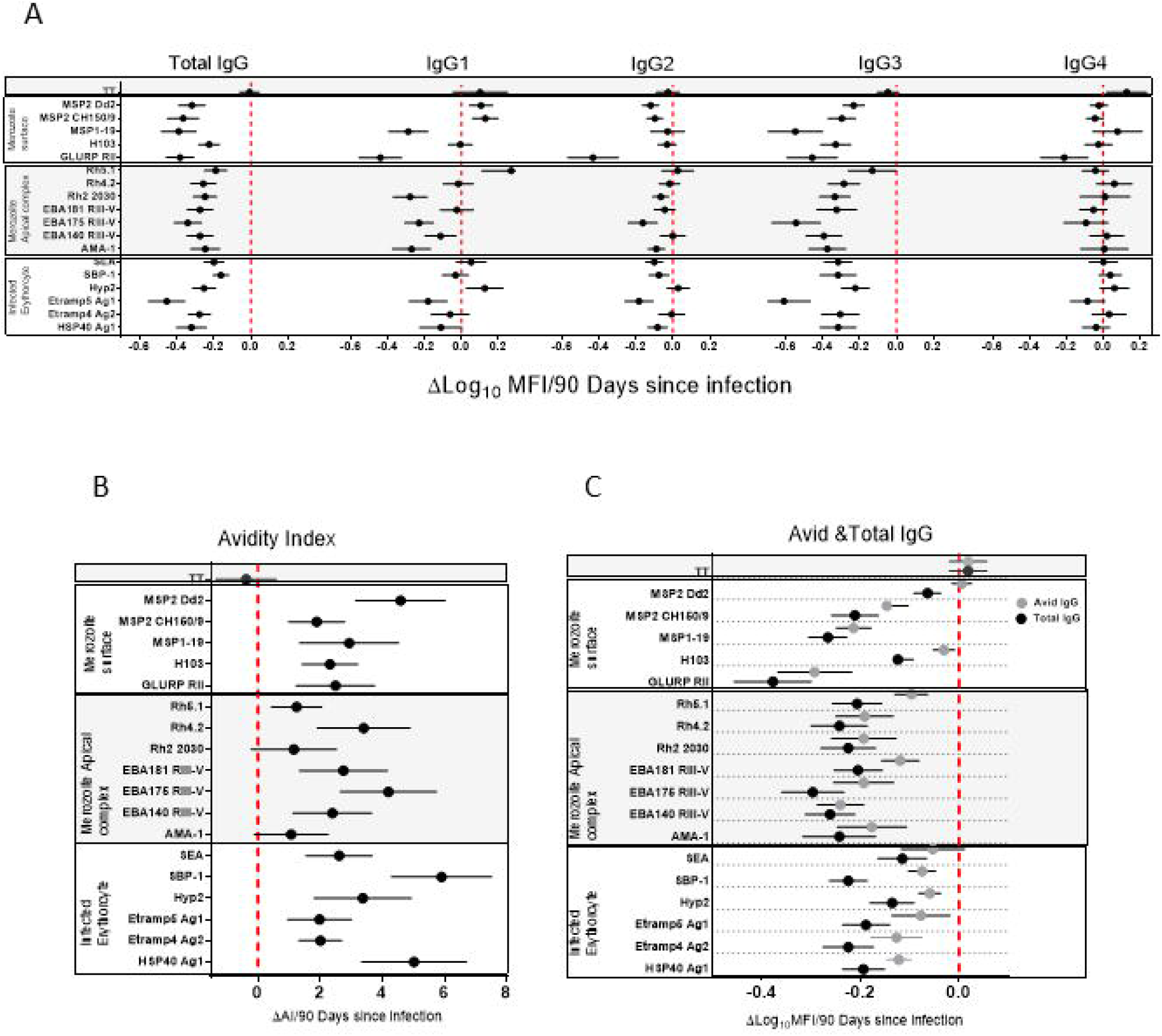
Association of total IgG, IgG1-4 (A), avidity index (B), avid and total IgG (C) 90 days following infection. The error bars represent mean and 95% confidence interval of the regression coefficient. Right of the zero red line is an increase and left is a decrease 90 days following infection. Overlap of the zero red line is a non-significant association.

### Increased *P. falciparum* specific antibody avidity index following infection was due to a preferential loss of low avidity antibodies

We hypothesized that the population level increase in antibody avidity index following a decrease in malaria transmission was due to the differential waning of total verses high avidity antibodies in the absence of blood stage infection. To test this hypothesis and measure the rate of decline in total verses avid antibodies, and the rate of change in the avidity index, we related antibodies and avidity index at all 4 time points (T1 - T4) to the duration of time since an individual’s last infection. Changes were estimated using log-linear or linear regression for antibody levels and avidity index respectively, i.e. assuming an exponential rate of change. The IgG avidity index increased following infection for all antigens except AMA-1 and Rh2_2030 (Figure 3B). The largest increase in avidity index following infection was seen for HSP40 ag1, SBP1 and MSP2_Dd2. There was no noticeable trend associated with the antigen categories of merozoite surface, merozoite apical complex and infected erythrocytes. As expected, the avidity index of IgG against tetanus toxoid did not change following *P. falciparum* infection, again confirming the specificity of the responses to the malarial antigens. In order to determine the driver of the increased avidity index in the absence of infection, the rate of total IgG decay was compared to avid IgG. The rate of decay following infection was higher in total compared to the avid IgG pool for all 18 antigens (Figure 3C). Because the avid IgG pool is a component of total IgG, the relative difference in the rate of decay following infection can be attributed to the non-avid pool. Thus, a preferential decay of non-avid antibodies is a likely explanation for the observed increase in avidity index.

### The rate of antibody decay following infection declined with age

Total IgG levels decayed faster in the younger (1 - 4 & 5 - 11yr) verses older (>18yr) age groups for the majority of the antigens (Figure 4A, C and Supplementary Figure 2). The relative difference in the decay slopes was more pronounced in the first 200 days following infection, and in IgG3 compared to IgG1. There was little change in the rate of decay with age for IgG2 and 4 for most antigens (Figure 4A, C and Supplementary Figure 2). The most pronounced changes across age category were observed for AMA-1, EBA175 RIII-V, EBA181 RIII-V, GLURP RII MSP2_Dd2 and MSP2_CH150/9. By contrast Etramp4 ag2, Etramp5 ag1 and HSP40 ag1 showed minimal changes across age. IgG1 and IgG3 had similar trends across the age categories for all the malaria antigens, with IgG3 showing more pronounced changes. The differences between ages were at maximum around 200 days following infection for total IgG, IgG1 and IgG3, driven by a rapid loss of antibodies associated with days since last infection in the 1 - 4 year age category compared to the older ages (5-11 & >18 yr).

**Figure 4.**
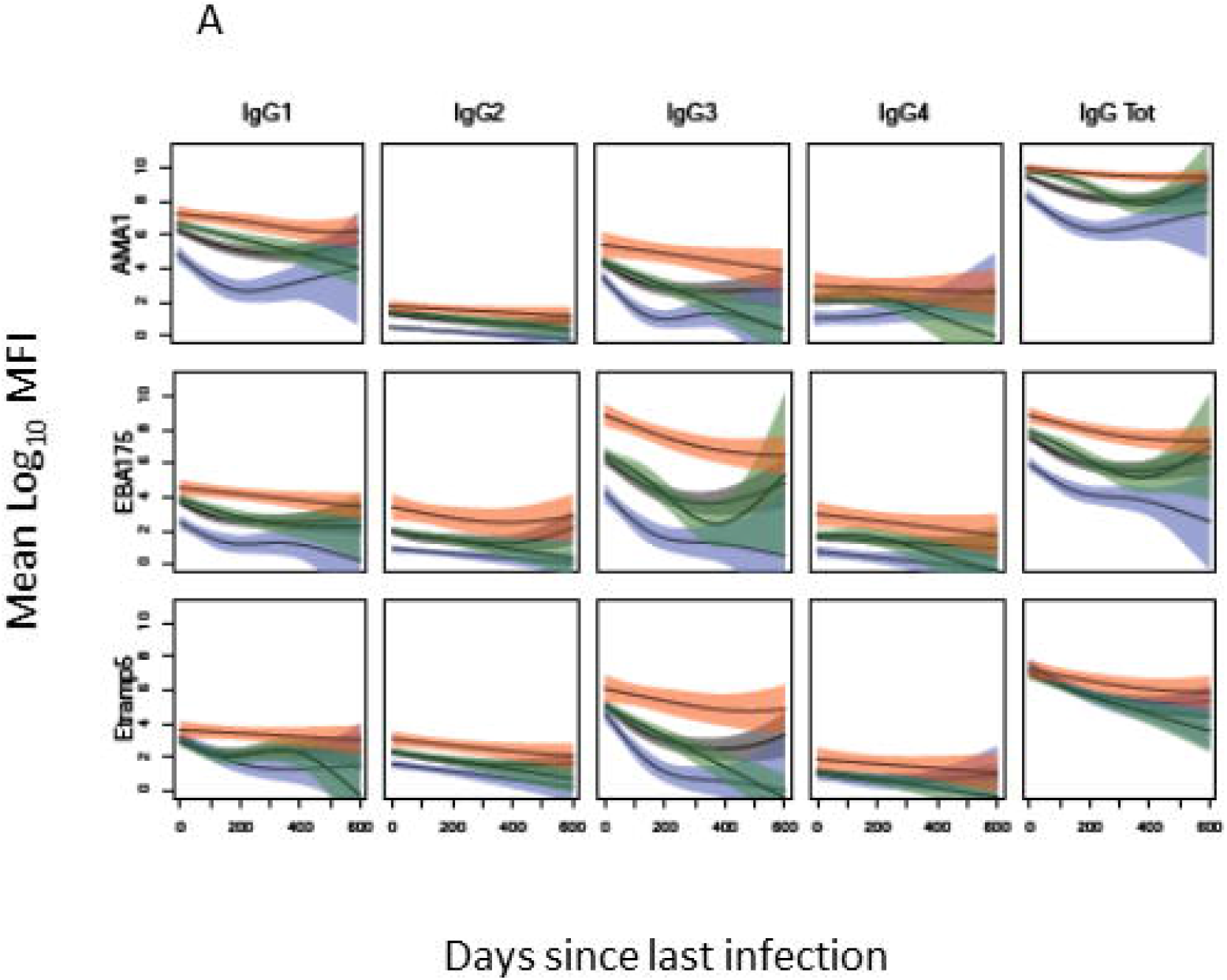
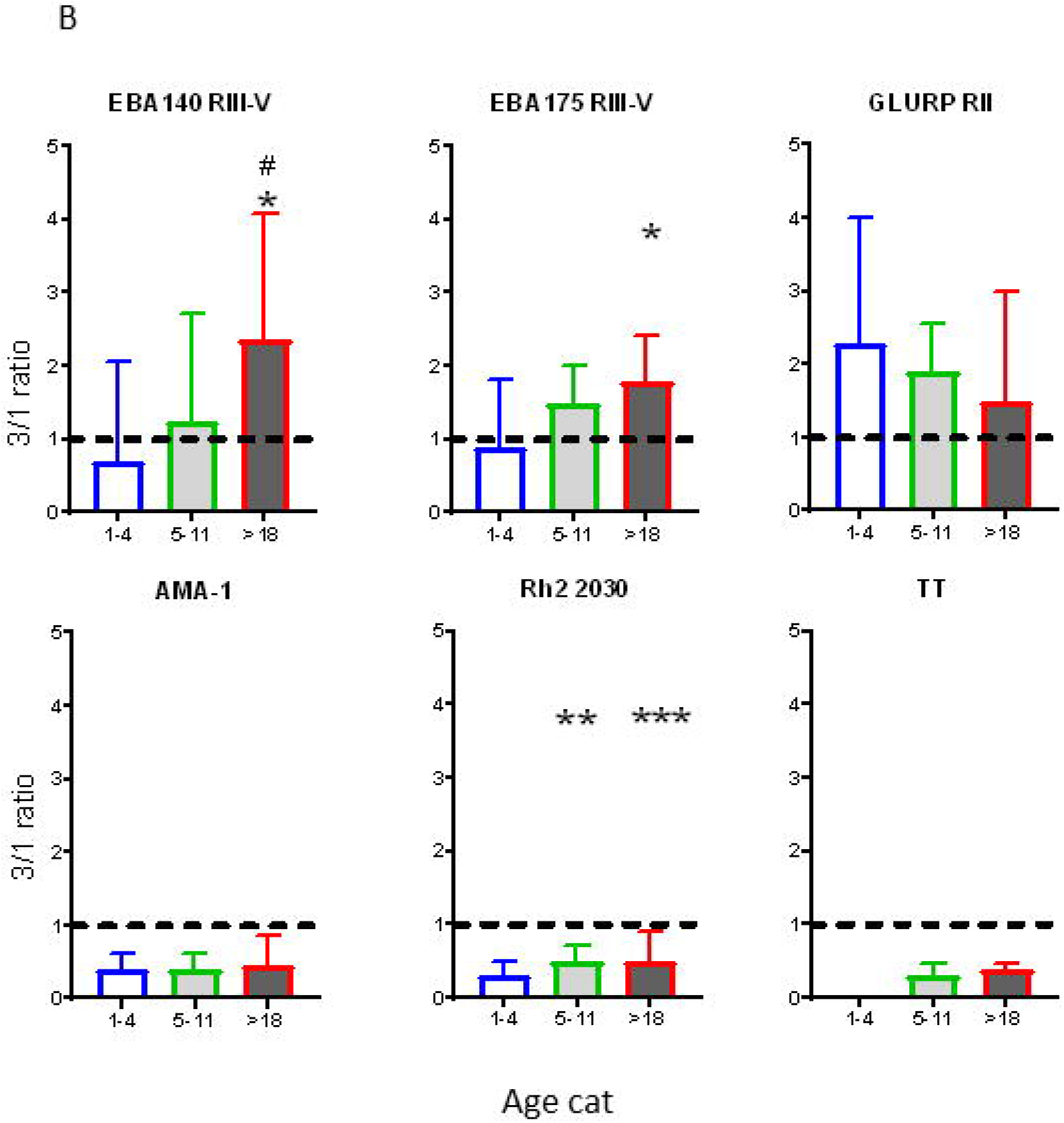
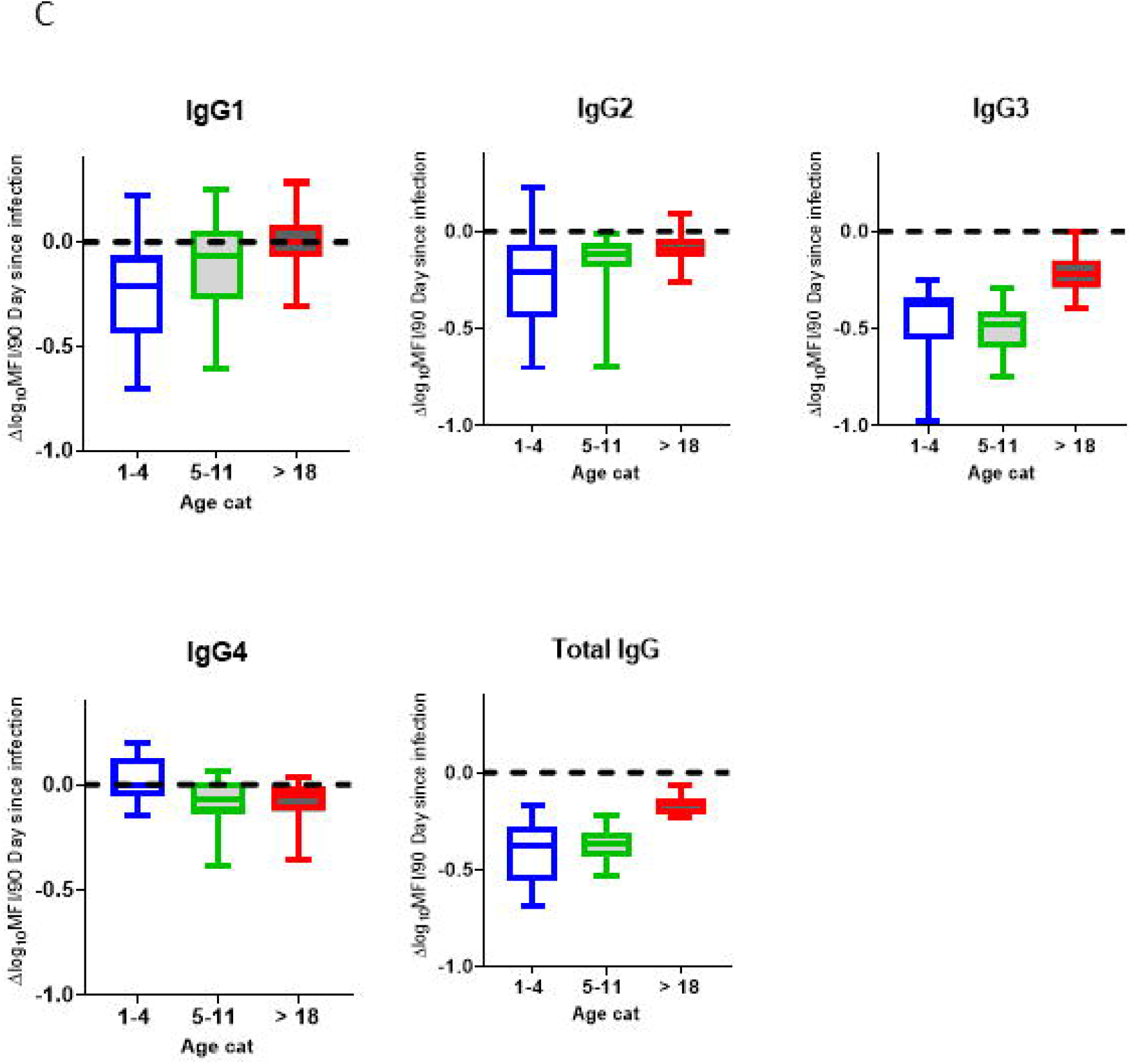
Changes in total IgG and subclasses 1 – 4 with age. (A) Representative plots of mean Log_10_ MFI against 90 days since last Infection, modeled using generalized additive models (GAMS), shaded areas represent 95% confidence interval. (B) Changes in IgG 3/1ratio with age. (C) Average rate (of 18 antigen) of decay following infection by age. Blue =1-4 years (N= 40), Green = 5-11 (N= 92) and Red = >18 years (N= 28). Gray = average (N=160) Ns = not significant, * = p= 0.05 – 0.01, ** p = 0.009 – 0.0001 *** p>0.00001.

To determine changes in the relative proportion of IgG1 and IgG3 with age, a ratio of Log_10_MFI IgG3 to IgG1 was determined and compared across the three age categories at T4 (to represent the antibody profile in the relative absence of recent infection). A ratio below 1 implied a higher proportion of IgG1 and a ratio above 1 a higher proportion of IgG3. For the 1 - 4 year age category the median IgG3/IgG1 ratio was below 1 for all antigens except for GLURP RII (Figure 4B and supplementary Figure 3). Median IgG3/IgG1 ratio increased for ages 5 - 11 and >18 years, approaching or exceeding 1 for most antigens except for AMA-1, Rh2_2030, HSP40 ag1 and Rh5.1, for which the median ratio remained below 1 in all age categories. By contrast, the IgG3/IgG1 ratio of tetanus toxoid remained below 1 for all age categories.

### High avidity antibodies increased but the avidity index decreased or remained unchanged with age

There was no association between age and avidity index for most of the antigens, except for Etramp4 ag2, EBA181 RIII-V, MSP2 Dd2 and AMA-1, for which avidity index was higher for age categories 1 - 4 compared with 5 - 11 and with >18 years (Figure 5). By contrast, there was an increase in high avidity antibodies across all age categories for all antigens except H103, particularly between age categories 5 - 11 and >18 years. There was a significant decrease in avidity index for TT between age categories 1 - 4 and 5 - 11 years, and an increase in the >18year age category. A similar pattern was observed for the high avidity antibodies. The restoration of both avidity index and high avidity antibodies may be attributed to a possible TT vaccination boost.

**Figure 5.**
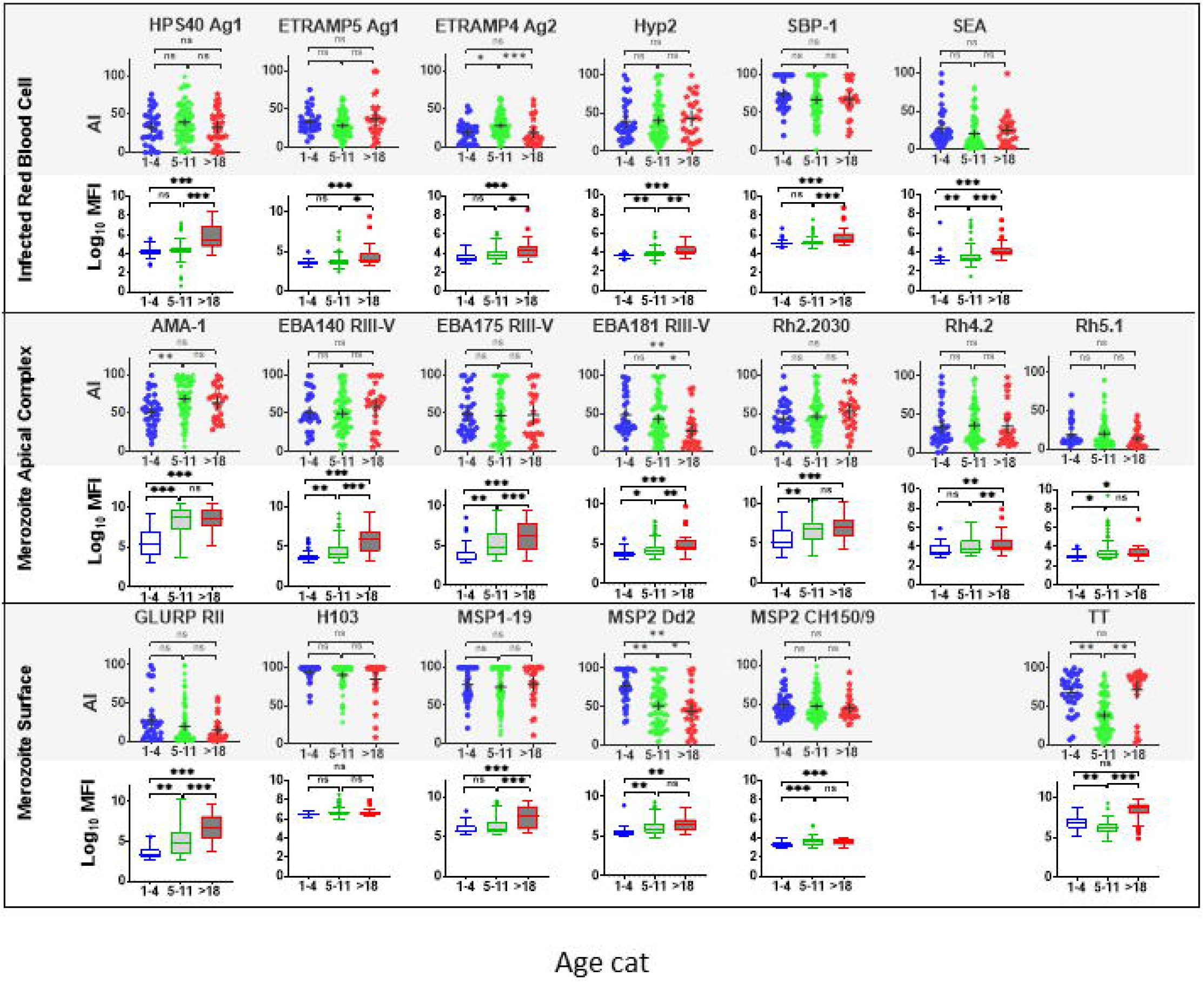
Comparison of Avidity index and high avidity antibodies across age category at T4. High avidity antibodies that remained binding after treatment with 2M GuHCl. Avidity index is the percentage of avid antibody of the total. Antibody was measure in a multiplex bead assay including 18 *P. falciparum* blood stage antigens and TT. Age categories at T4 include 1-4 years (N= 40), 5-11 (N= 92) and >18 years (N= 28). Ns = not significant, * = p= 0.05 – 0.01, ** p = 0.009 – 0.0001 *** p>0.00001

## Discussion

We measured responses to a panel of 18 *P. falciparum* blood stage antigens to gain broad insight into the dynamics of antibody levels and avidity when malaria transmission is reduced. Consistent with previous studies, we showed that total IgG increased with age (46), that it substantially declined when exposure decreased, that IgG1 and IgG3 were the dominant subclasses, and that there was antigen specific bias of dominance to these subclasses (20,21,25). In addition to previous findings, we showed that the decrease in total IgG was consistently driven by the decrease in IgG3 and occasionally by a decrease in IgG1. We also showed that the proportion of IgG3 relative to IgG1 maintained in the absence of infection increased with age, and that the rate of antibody decay following infection declined with age.

We validated previous findings that the antimalarial antibody avidity index increased following infection and was not consistently associated with age (39,50). In addition to previous work (47), we showed that the increase in avidity index following infection was likely due to the more rapid loss of the low avidity vs. high avidity antibodies.

Differences in individual antibody avidity is likely to be on a continuous spectrum. Use of GuHCl at one concentration to define avidity results in dichotomization of the data. Therefore, though useful, the designation of ‘avid’ and ‘non-avid’ is an over-simplification. We postulate that the low avidity antibodies are produced mainly by short-lived plasma cells (SLPC), and high avidity antibodies by long lived plasma cells (LLPC). Infection drives expansion of the SLPC and non-avid antibodies, and the rapid contraction of SLPCs following infection leads to rapid preferential decay of the low avidity antibodies. Therefore, the increase in avidity index we observed may indicate affinity maturation per se and/or the relative contributions of SLPCs and LLPCs following infection. A study conducted in the setting of ongoing malaria transmission and infection in Nigeria found that antibody avidity was stable among most individuals over time (56); this finding is largely consistent with our proposed model. In future studies, the direct quantification of high avidity antibodies - not performed here - may be a better indicator of affinity maturation compared to the avidity index. For example, we demonstrated that high avidity antibodies increased with age despite the lack of an increase in the avidity index with age.

The implication of early acquisition of the predominantly IgG1 response we observed is not well understood, however there is increasing evidence to suggest that IgG3 may be more protective than other subtypes against malaria. A stronger association between IgG3 and a reduced risk of malaria compared to IgG1 was reported against PfRH5 and PfRipr,(27). In addition IgG3 C1q fixing antibodies to MSP2 and EBA175 RIII-V were associated with parasite clearance following treatment (57). IgG3 has greater activity in promoting complement fixation and opsonic phagocytosis, which are mechanisms that appear to contribute to protective immunity (21,23,58). The observation that IgG3 decayed at a faster rate than IgG1 in the absence of infection, and was significantly slower in adults compared to infants, implies a gradual accumulation of IgG3^+^ LLPC with age and exposure.

The finding of a shift in dominance from IgG1 to IgG3 with age is intriguing, especially when it is considered that subclass switching generally proceeds in a single direction (IgG3, to IgG1, to IgG2 and IgG4) due to the nature of the CH domains in the Fc region (5). Therefore, memory B cells (MBC) that eventually give rise to IgG3^+^ LLPC cannot be the cells that switched to IgG1. Rather, naïve B cells, IgG3^+^ MBC, or IgM^+^ MBC are likely to be the source of IgG3^+^LLPC that gradually accumulate with age (59). Some studies have not found a major effect of age and exposure on the IgG subclass responses to antigens (33) suggesting the nature of exposure or specific epidemiological conditions may influence this effect.

IgG3 has the shortest intrinsic half-life, at 7 days, compared to ~21 days for the other subclasses (60). This implies that maintaining a large pool of IgG3 in the absence of infection requires a larger pool of LLPC compared to IgG1. We observed a slower decay or increased half-life with age of IgG1 and IgG3, but with a shift in relative proportion from predominantly IgG1 to IgG3. This may imply gradual expansion of preferentially IgG3^+^ LLPC with age. Previous studies have indicated a requirement of high antibody titers to maintain immunity to malaria (15). As IgG3 is the predominant IgG for most of the malaria antigens we evaluated, a larger pool of IgG3^+^ LLPC may be required to maintain high titers compared to IgG1 in the absence of infection. While IgG3 is suspected to be functionally more potent compared to IgG1 (61), being the dominant subclass poses a limitation of poor maintenance of protective levels in the absence of infection. This limitation poses a big challenge to malaria vaccine development, especially in endemic areas where existing memory B cells are inclined to switch to IgG3^+^ LLPC. A recent analysis of RTS,S vaccine responses in children found IgG3 responses had a higher decay rate than IgG1(62). Antigens such as AMA-1 and Rh2_2030 elicited a predominantly IgG1 response across the age groups. In addition, these antigens had a relatively lower decay rate following infection compared to antigens that were predominantly IgG3. This observation implies that some *P. falciparum* antigens may have inherent features that influence IgG class switch bias. Therefore, we need to better understand the functional potency of IgG1 verses IgG3 and antigen features that lead to biased class switching to inform malaria vaccine designs.

## Conclusions

In the setting of declining exposure to *P. falciparum*, IgG3 was the major driver of observed antibody decay rate and avidity index increased, likely due to preferential decay of the non-avid vs. avid antibodies. More direct quantification of avid antibodies, ascertainment of the cellular phenotype producing various antibodies over time, and functional studies of various aspects of antibody quality will be necessary to understand their protective roles in malarial immunity. This knowledge is critical for development and assessment of vaccine strategies that can improve longevity of malaria vaccine efficacy and in understanding the longevity of immunity to malaria as transmission declines.

## Supplemental information

### Figure legend

**Figure S1.** Dot Plot of log_10_ MFI of Merozoite apical complex. Antibody levels were measured using a MagPix (Luminex) multiplex assay including 18 malaria blood stage antigens and TT. Each dot represents a single sample from 160 participants at 4 time points. The horizontal bar represents the median and 95% CI.

**Figure S2.** Changes in total IgG and subclasses 1 – 4 with age. Total IgG and IgG1 – 4 for 19 *P falciparum* antigens and TT was measure in a multiplex bead array assay. Plots of mean Log_10_ MFI against days since last Infection, modeled using generalized additive models (GAMS), shaded areas represent 95% confidence interval.

**Figure S3.** Changes in total IgG3/IgG1 ratio with age. Total IgG and IgG1 – 4 for 19 *P falciparum* antigens and TT was measure in a multiplex bead array assay. Blue =1-4 years (N= 40), Green = 5-11 (N=92) and Red = >18 years (N= 28). Gray = average (N=160) Ns = not significant, * = p= 0.05 – 0.01, ** p = 0.009 – 0.0001 *** p>0.0000.

### Table

**Table S1** Summary of the *P.falciparum* Blood stage antigens

