## Supplementary figures and images for "Impact of a rapid decline in malaria transmission on antimalarial IgG subclasses and avidity"

### Figure S1

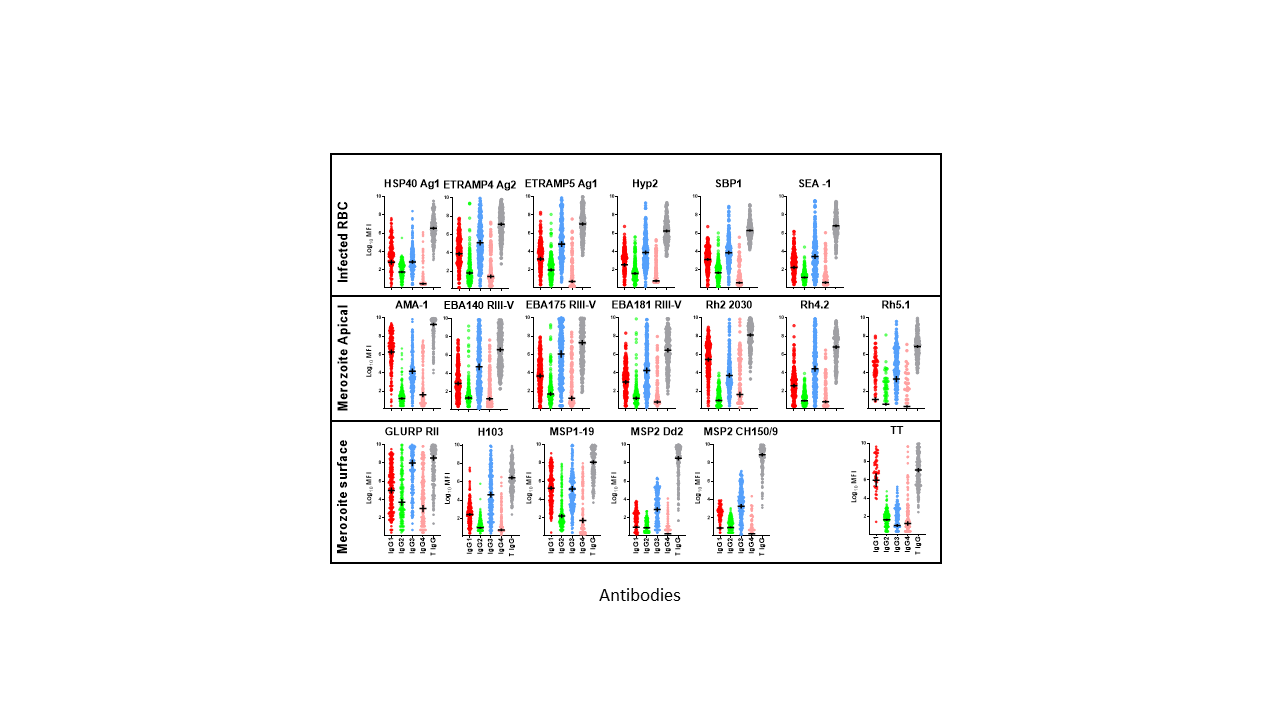

### Figure S2

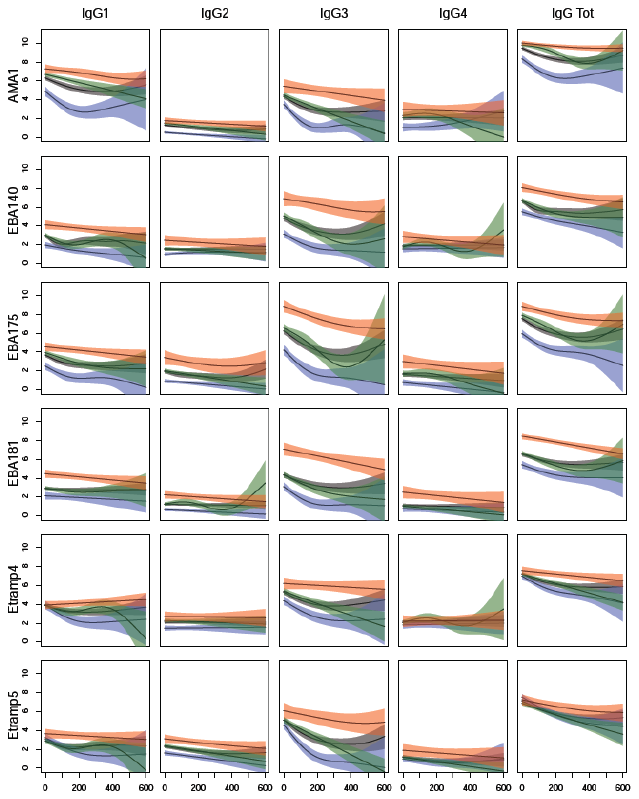

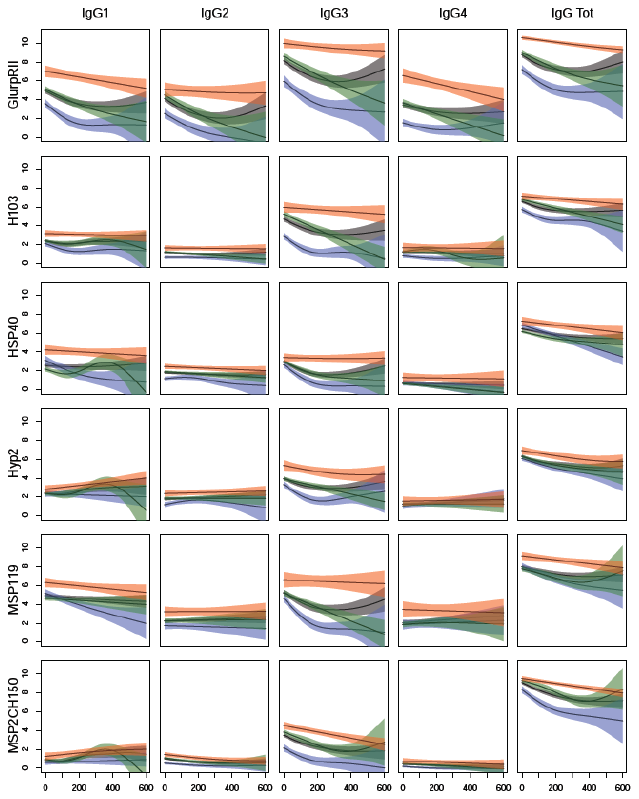

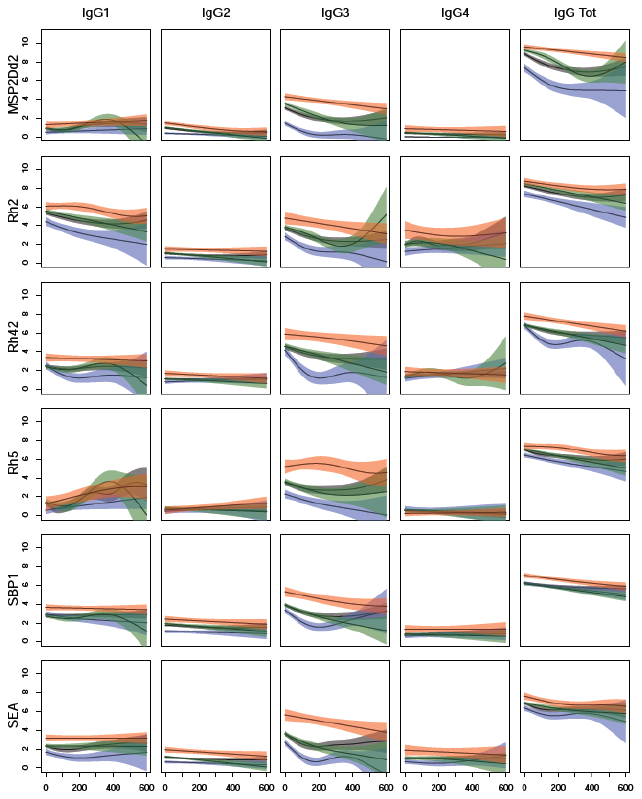

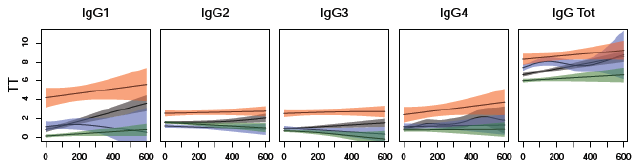

### Figure S3

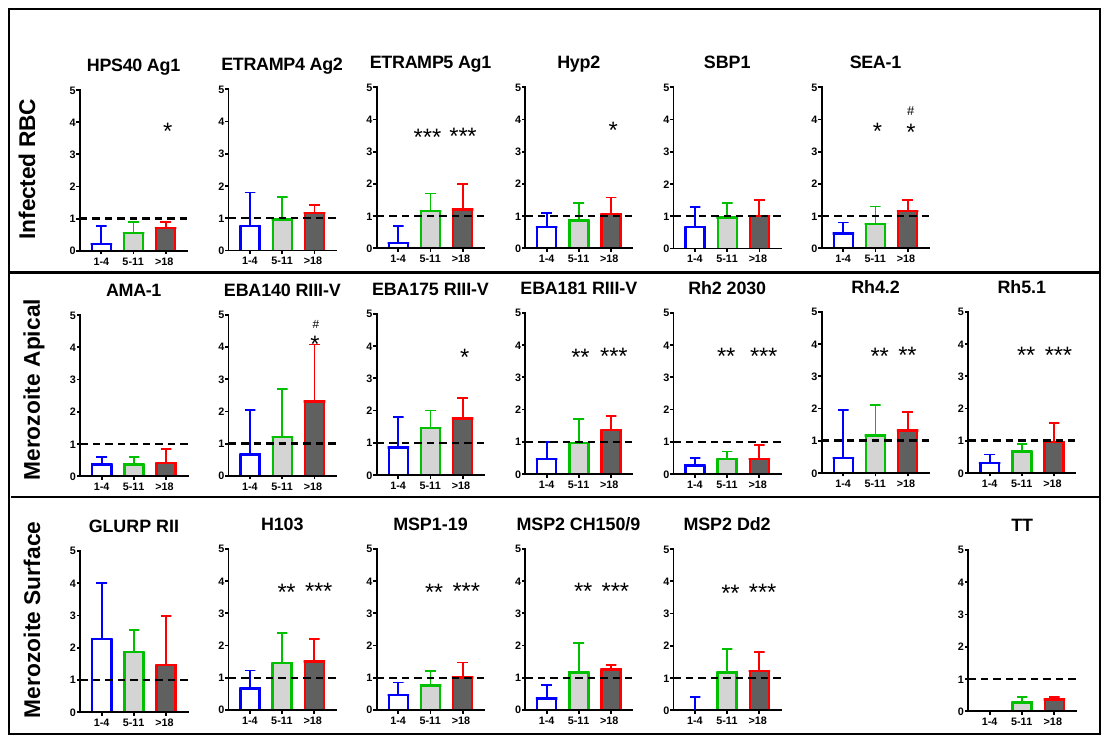
