## Supplementary material for "Impact of a rapid decline in malaria transmission on antimalarial IgG subclasses and avidity": Table S1

| Gene ID | Description | Antigen name | Allele | AA | Location | Tag |
|---|---|---|---|---|---|---|
| PF3D7_0501100.1 | Heat Shock Protein 40, type II, Antigen1(*KT) | HSP40 ag1 | 3D7 | 71-153 | iRBC/Gam | GST |
| PF3D7_0423700 | Early Transcribed Membrane Protein 4, Antigen2 (*KT) | Etramp4 ag2 | 3D7 | 76-137 | iRBC/PVM | GST |
| PF3D7_0532100 | early transcribed membrane protein 5 Antigen1(*KT) | Etramp5 ag1 | 3D7 | 26-111 | iRBC/PVM | GST |
| PF3D7_1002000 | Plasmodium exported protein (hyp2) unknown function (*KT) | Hyp2 | 3D7 | 101-418 | iRBC/PVM | GST |
| PF3D7_0501300 | Skeleton-Binding Protein 1(*KT) | SBP1 | 3D7 | 1-239 | Scht/MC | GST |
| PF3D7_1021800 | Schizont Egress Antigen 1(*KT) | SEA-1 | 3D7 | 810-1083 | Scht/MC | GST |
| PF3D7_1133400 | Apical Membrane Antigen 1(63) | AMA-1 | FVO | 97-546 | SpZ/Mer | $His_{x6}$ |
| PF3D7_1301600 | Erythrocyte Binding Antigen-140 Region III-V (34) | EBA140 RIII-V | 3D7 | | Mer -M | GST |
| PF3D7_0731500 | Erythrocyte Binding Antigen-175 Region III-V | EBA175 RIII-V | 3D7 | 761-1298 | Mer-M | GST |
| PF3D7_0102500 | Erythrocyte Binding Antigen-181 Region III-V (34) | EBA181 RIII-V | 3D7 | 769-1365 | Mer -M | GST |
| PF3D7_1335400 | Reticulocyte Binding Protein Homologue 2 (64) | Rh2 | D10 | 2030-2528 | Mer-Rh | GST |
| PF3D7_0424200 | Reticulocyte Binding Protein Homologue 4 (65) | Rh4.2 | 3D7 | 28-766 | Mer-Rh | $His_{x6}$ |
| PF3D7_0424100 | reticulocyte binding protein | Rh5 | 3D7 | 26-526 | Mer-Rh | C- |
| PF3D7_1035300 | Glutamate Rich Protein R2 (67) | GLURP RII | F32 | 816-1091 | Mer-S | n/a |
| PF3D7_1036000 | Merozoite Surface Protein 11/H101 (68) | H103 | 3D7 | 40-243 | Mer-S | GST |
| PF3D7_0930300 | 19kDa fragment of MSP1 molecule (69) | MSP1_19 | Wellcome | 1631-1726 | Mer-S | GST |
| PF3D7_0206800 | Merozoite surface protein 2, Dd2 allele (70) | MSP2 Dd2 | Dd2 | 22-247 | Mer-S | GST |
| PF3D7_0206800 | Merozoite surface protein 2, CH150/9 allele (70) | MSP2 CH150/9 | CH150/9 | 34-215 | Mer-S | GST |
| n/a | Tetanus Toxoid (Non-adsorbed) | TT | n/a | n/a | n/a | n/a |

iRBC =Infected red blood cell, Gem- Gametocyte, PVM = parasitophorous vacuole membrane, MC = Maurer's cleft, SPZ = sporozoite, Mer-S = merozoite surface, Mer-M = merozoite micronemes, Mer-Rh = merozoite Rhoptry   *KT = Tetteh. K, unpublished
